# Oligomerization-driven MLKL ubiquitylation antagonises necroptosis

**DOI:** 10.1101/2021.05.01.442209

**Authors:** Zikou Liu, Laura F. Dagley, Kristy Shield-Artin, Samuel N. Young, Aleksandra Bankovacki, Xiangyi Wang, Michelle Tang, Jason Howitt, Che A. Stafford, Ueli Nachbur, Cheree Fitzgibbon, Sarah E. Garnish, Andrew I. Webb, David Komander, James M. Murphy, Joanne M. Hildebrand, John Silke

## Abstract

Mixed lineage kinase domain-like (MLKL) is the executioner in the caspase-independent form of programmed cell death called necroptosis. Receptor Interacting serine/threonine Protein Kinase 3 (RIPK3) phosphorylates MLKL, triggering MLKL oligomerization, membrane translocation and membrane disruption. MLKL also undergoes ubiquitylation during necroptosis, yet neither the mechanism nor significance of this event have been demonstrated. Here we show that necroptosis-specific, multi-mono-ubiquitylation of MLKL occurs following its activation and oligomerization. Ubiquitylated MLKL accumulates in a digitonin insoluble cell fraction comprising plasma/organellar membranes and protein aggregates. This ubiquitylated form is diminished by a plasma membrane located deubiquitylating enzyme. MLKL is ubiquitylated on at least 4 separate lysine residues once oligomerized, and this correlates with proteasome- and lysosome-dependent turnover. Using a MLKL-DUB fusion strategy, we show that constitutive removal of ubiquitin from MLKL licences MLKL auto-activity independent of necroptosis signalling in mouse and human cells. Therefore, besides its role in the kinetic regulation of MLKL-induced death following an exogenous necroptotic stimulus, ubiquitylation also contributes to the restraint of basal levels of activated MLKL to avoid errant cell death.

## Introduction

Necroptosis is a type of programmed cell death that shares some molecular components with the better-known apoptotic cell death pathway, but has distinct morphological features and different physiological consequences. Unlike apoptosis, which is immunologically silent and can be rapidly cleared by neighbouring phagocytic cells (Segawa & Nagata, 2015), necroptosis induces an inflammatory response by releasing cellular contents including DNA and cytosolic proteins (Kaczmarek *et al*, 2013). Necroptosis can be induced by a number of stimuli, but is mainly studied downstream of Tumor Necrosis Factor (TNF) ligation to its receptor Tumor Necrosis Factor Receptor 1 (TNFR1). If Inhibitor of APoptosis (IAP) proteins and caspase activities are suppressed, this results in higher order assemblies of Receptor Interacting serine/threonine Protein Kinase 1 (RIPK1) and RIPK3, and subsequent RIPK3 activation and auto-phosphorylation (Sun *et al*, 2002; Wu *et al*, 2014). MLKL is phosphorylated by RIPK3 and oligomerizes, translocates to biological membranes and induces organelle and cell swelling and membrane rupture (Petrie *et al*, 2019a; Vanden Berghe *et al*, 2015).

Ubiquitylation plays a pivotal role in regulating TNF signalling. In addition to RIPK1 ubiquitylation by cIAPs, which forms a platform for MAPK and NF-κB activation (Bertrand *et al*, 2008), the Linear Ubiquitin Chain Assembly Complex (LUBAC), composed of HOIL-1, HOIP and SHARPIN, generates M1-linked ubiquitin chains on RIPK1, TRADD, TNFR1 and NEMO. This ubiquitylation leads to full activation of IKK and MAPK, and limits TNF induced cell death (Haas *et al*, 2009; Ikeda *et al*, 2011; Tokunaga *et al*, 2009). LUBAC also recruits CYLD, a deubiquitylating enzyme (DUB), that removes M1- and K63-linked ubiquitin chains on RIPK1 and other complex components, while A20 sterically protects M1-linked ubiquitin chains (Dondelinger *et al*, 2016; Draber *et al*, 2015).

Ubiquitylation has been implicated in the regulation of signalling checkpoints during necroptosis. RIPK3 undergoes K63-linked ubiquitylation on Lys-5, which is believed to support the formation of necrosome. Removal of this ubiquitin chain by the ubiquitin-editing enzyme A20 was proposed to negate necroptosis, because A20 loss led to RIPK3-dependent necroptosis in T cells and fibroblasts (Onizawa *et al*, 2015). Furthermore, ubiquitylated TRAF2 was reported to be associated with inactive MLKL, and CYLD deubiquitylates TRAF2 following necroptotic stimulation, allowing TRAF2 to dissociate from MLKL and MLKL to engage with, and be activated by, RIPK3 (Petersen *et al*, 2015).

MLKL has also been shown to undergo ubiquitylation upon necroptotic stimulation (Hildebrand *et al*, 2020; Lawlor *et al*, 2015), yet the significance of this post-translational modification and its signalling function are unknown. Here we show that necroptotic signalling stimulates MLKL ubiquitylation and that this antagonises necroptosis *via* restraining the protein level of activated MLKL. MLKL oligomerization is the crucial stage for MLKL ubiquitylation, but RIPK3 phosphorylation is not necessary because auto-active oligomerizable MLKL mutants are ubiquitylated. MLKL ubiquitylation was removed *in vitro* by USP21, but not by other chain specific deubiquitylating enzymes, suggesting that MLKL is mono-ubiquitylated at multiples sites. Upon necroptotic stimulation, MLKL variants that are unable to induce cell death can still be ubiquitylated and this correlates with proteasome/lysosome mediated turnover. Conversely, mutation of four lysine residues identified by mass spectrometry analysis to be ubiquitylated following a necroptotic stimulus did not affect MLKL ubiquitylation or MLKL’s cytotoxic activity. We therefore devised a novel approach to completely remove all ubiquitins from MLKL by fusing it to USP21. We showed that this MLKL-USP21 fusion was resistant to ubiquitylation in both human and mouse cell lines and was more cytotoxic when compared with MLKL fused to a catalytically-inactive USP21. Strikingly, human MLKL, which is very resistant to auto-activation (Petrie *et al*, 2018; Tanzer *et al*, 2016), was auto-activated by the USP21 fusion suggesting that ubiquitylation is a major break on the activation of human MLKL.

## Results

### MLKL becomes ubiquitylated during necroptosis

MLKL has previously been reported to undergo ubiquitylation in mouse bone marrow-derived macrophages (BMDMs) stimulated with the necroptotic stimulus: LPS/Smac-mimetic (Compound A)/pan-caspase inhibitor (Q-VD-OPh) (Lawlor *et al.,* 2015). To test whether MLKL ubiquitylation is specific to necroptosis, we stimulated wildtype mouse dermal fibroblasts (MDFs) with TNF (T), Smac-mimetic (S) and the pan-caspase inhibitor IDN-6556 (I) individually or in combination for 3 hrs. GST-UBA immobilised on glutathione sepharose beads (Hospenthal *et al*, 2015; Stafford *et al*, 2018) was used to purify ubiquitylated proteins from cell lysates. These purified samples generated a distinct ladder of between 50 and 75 kDa when probed with anti-MLKL, which corresponds to non-ubiquitylated MLKL, mono-ubiquitylated MLKL and multi-ubiquitylated MLKL (**Fig. 1A**). Ubiquitylated MLKL species were only clearly detected for stimuli that promote MLKL phosphorylation (TI and TSI), but not for stimuli that induce apoptotic cell death (TS) (**Fig. 1A**). Ubiquitylation appears to occur in the same time window as membrane permeabilization, because by the time of UBA-pulldown (UBA-PD) for MDFs (3 hrs) and HT29 cells (16 hrs), cells were ~60% propidium iodide (PI)-positive (**Supp. Fig. 1A**).

**Figure 1.**
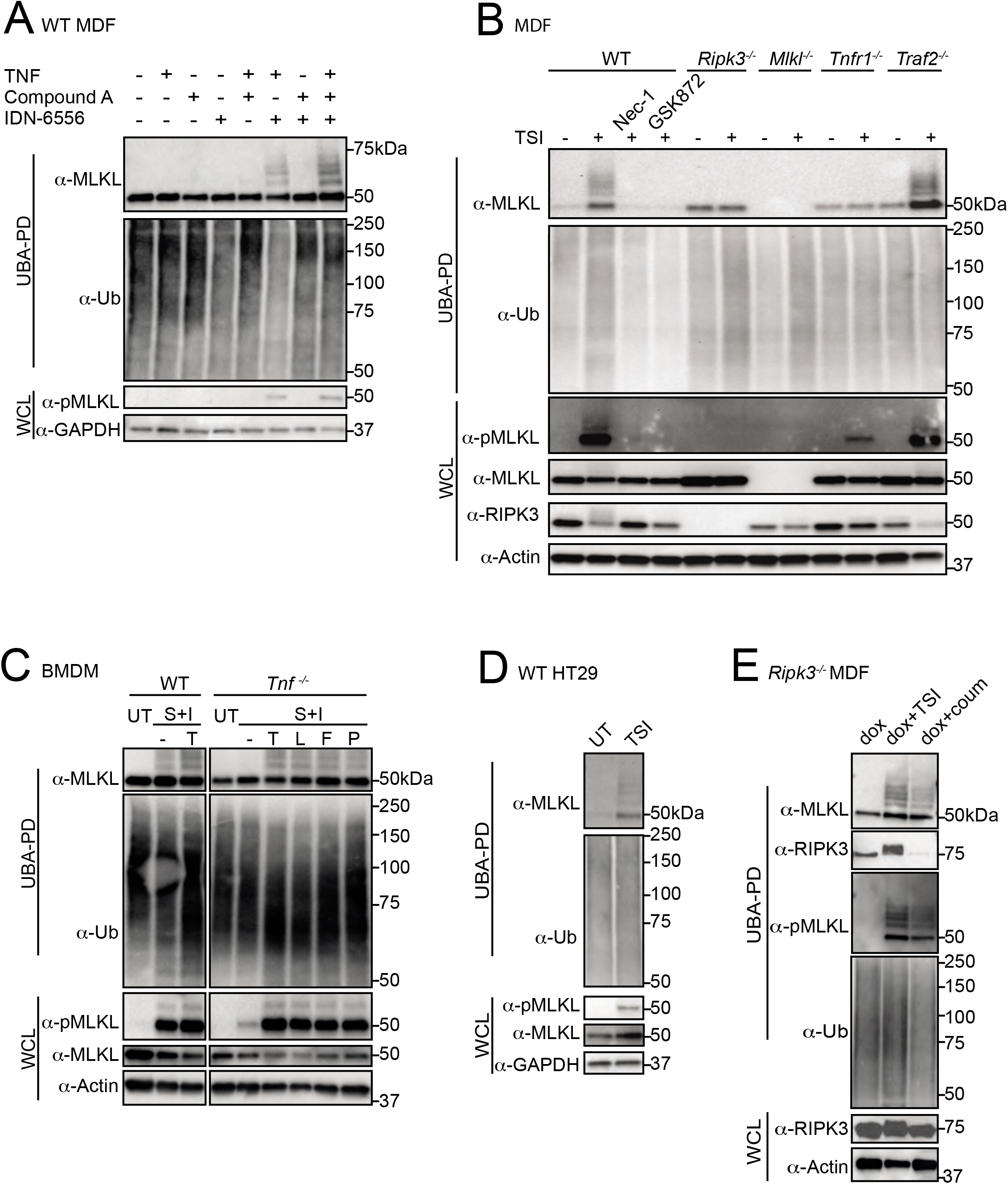
MLKL undergoes ubiquitylation during necroptosis. A WT MDFs were treated ± TSI individually or in combination for 3 hrs. Whole cell lysates (WCL) and UBA-pull down (UBA-PD) fractions were analysed by Western blot and probed with antibodies as indicated. Representative of three independent experiments. Samples of UBA-pull down in following figures were analysed in the same way unless otherwise indicated. B WT, *Ripk3^-/-^, M1k1^-/-^, Tnfr1^-/-^* and *Traf2^-/-^* MDFs were untreated (UT) or treated with TSI for 3 hrs. Nec-1 and GSK872 were added to inhibit RIPK1 and RIPK3 kinase activities respectively. C WT and *Tnf^-/-^* BMDMs were treated ± death ligands including TNF (T), LPS (L), Fas ligand (F) and Poly I:C (P) in addition to S and I for 3 hrs. D WT HT29 cells were untreated (UT) or treated with TSI for 16 hrs. E RIPK3-gyrase were inducibly expressed in *Ripk3^-/-^* MDFs by doxycycline (dox) for 5 hrs, and cells were then treated ± combination of TSI, or coumermycin (coum) for 3 hrs.

As expected, TSI failed to induce MLKL ubiquitylation in *Tnfr1^-/-^* MDFs, but, interestingly, did not completely prevent all TSI induced MLKL phosphorylation (**Fig. 1B**). Furthermore, the kinase activities of RIPK1 and RIPK3 were required for MLKL ubiquitylation because genetic deletion of *Ripk3,* or treatment with the RIPK1 and RIPK3 inhibitors, Nec-1 and GSK872, respectively, prevented MLKL ubiquitylation (**Fig. 1B**). MLKL phosphorylation and ubiquitylation were not, however, reduced by genetic deletion of *Traf2* (**Fig. 1B**). To test whether MLKL ubiquitylation occurred in other cell types, we examined mouse BMDMs. Necroptosis can be induced in BMDMs by a range of different ligands including LPS, Poly I:C and the TNF-receptor superfamily ligand FasL, when they are combined with a Smac-mimetic and a caspase inhibitor, IDN-6556 (SI) (Holler *et al*, 2000; Kaiser *et al*, 2013). To exclude a role for autocrine TNF, whose synthesis can be induced by a Smac-mimetic, LPS, poly I:C or FasL, we compared BMDMs isolated from *Tnf^-/-^* and WT mice, and saw the same MLKL ubiquitin laddering pattern in both. This ubiquitylation correlated with MLKL phosphorylation on S345, a well-known hallmark of MLKL activation and necroptosis (**Fig. 1C**) (Murphy *et al*, 2013). We and others have shown that human and mouse MLKL are regulated distinctly (Davies *et al*, 2020; Davies *et al*, 2018; Li *et al*, 2015; Petrie *et al.,* 2018; Tanzer *et al.,* 2016), therefore to see whether human MLKL was also ubiquitylated during necroptosis, we treated the human colonic adenocarcinoma HT29 cell line with TSI. Although induction of necroptosis takes longer in HT29 than in MDFs (**Supp. Fig. 1B**), we observed similar MLKL phosphorylation and ubiquitylation in HT29 cells as in MDFs with similar levels of cell death (**Fig. 1D, Supp. Fig. 1B**).

To determine to what extent upstream necroptotic signalling was required for MLKL ubiquitylation, we used a well-established RIPK3 dimerization strategy to activate RIPK3 in the absence of necroptotic signals (Moujalled *et al*, 2014; Orozco *et al*, 2014). We therefore stably expressed a doxycycline-inducible RIPK3-gyrase fusion protein, which can be dimerized and activated by coumermycin (Moujalled *et al.,* 2014), in *Ripk3^-/-^* MDFs. As expected, forced dimerization of RIPK3 induced MLKL phosphorylation but was also able to induce MLKL ubiquitylation independent of TSI stimulation and to similar levels as TSI, indicating that RIPK3 dimerization and activation is sufficient to induce MLKL ubiquitylation (**Fig. 1E**).

### MLKL is ubiquitylated on multiple sites during necroptosis

Ubiquitin can be coupled to, predominantly, lysine residues in target proteins. Ubiquitin itself contains 8 ubiquitylation sites (M1, K6, K11, K27, K29, K33, K48 and K63), which leads to the formation of ubiquitin chains of varying architectures. Differently linked Ub chains serve distinct functions in cells (Swatek & Komander, 2016). To investigate the ubiquitin architecture on MLKL, we purified ubiquitylated proteins from necroptotic cells using sepharose beads coupled with GST-UBA and digested them on the beads with a set of linkage selective deubiquitylating enzymes (UbiCRest) (Hospenthal *et al.,* 2015). Each DUB specifically recognizes and removes a known subset of poly-ubiquitin chain type with validated activity (**Fig. 2A**) (Stafford *et al.,* 2018). The non-specific DUB USP21 (Ye *et al*, 2011) converted MLKL into a non-ubiquitylated form, but the other chain-specific DUBs made no discernible difference to the MLKL ubiquitin laddering pattern (**Fig. 2B**). To control that there was sufficient DUB activity we decreased the ratio of substrate to DUBs to 1/2 and 1/4 (**Fig. 2C**). vOTU, a DUB that can cleave every type of poly-ubiquitin chain except M1-linked chains, deubiquitylated most ubiquitylated proteins (see total ubiquitin blot, **Fig. 2C**), yet it only deubiquitylated the highest molecular weight species of MLKL~Ub adducts, and did not affect bands representing the lower molecular weight MLKL~Ub species. No changes were observed upon treatment with OTULIN, a DUB specific for M1-linked ubiquitin chains (Keusekotten *et al*, 2013), indicating that the residual Ub left by vOTU are not M1-linked ubiquitin chains. Given that USP21 is the only DUB in the tested panel that can cleave the covalent bond between a protein substrate and the first ubiquitin unit added, these data suggest that ubiquitylated MLKL is most likely mono-ubiquitylated at multiple sites.

**Figure 2.**
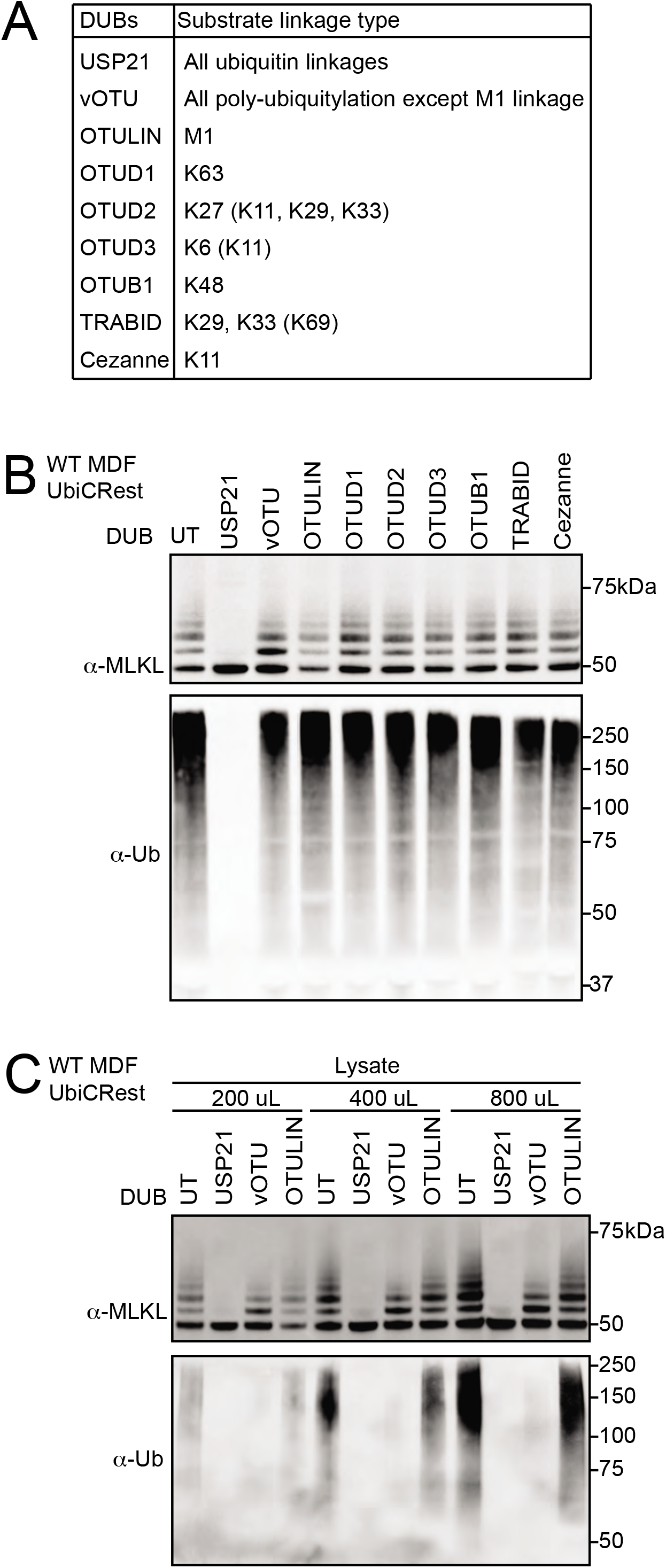
MLKL is mono-ubiquitylated at multiple sites. A Deubiquitylating enzymes (DUBs) and their ubiquitin substrates. Less efficiently cleaved substrates are indicated in brackets. B UBA-pull down from WT MDFs treated with TSI for 3 hrs were subjected to the DUBs shown in (A). Beads eluates were analysed by Western blot and probed with antibodies as indicated. Representative of three independent experiments. C 1.4 mL cleared cell lysate from 4 x 10^6^ TSI-treated WT MDFs was split into three parts of the indicated volume, followed by UBA-pull down and DUB incubation. Bead eluates were analysed by Western blot and probed with the indicated antibodies. Representative of three independent experiments.

### MLKL ubiquitylation accumulates in the membrane fraction and can be deubiquitylated by USP21 localised to the plasma membrane

During necroptosis, activated MLKL oligomerizes and translocates to biological membranes, ultimately causing membrane rupture and cell death (Cai *et al*, 2014; Chen *et al*, 2014; Hildebrand *et al*, 2014). To understand more about MLKL ubiquitylation, we examined when and where this occurs. WT MDFs were stimulated with TSI over a time course from 30 to 180 minutes. Cytosolic and crude membrane fractions were generated as previously described using digitonin (Liu *et al*, 2018), and subjected to UBA-pull down. We refer to the 0.025% digitonin insoluble fraction as ‘crude membrane’, but do not exclude the potential for non-membrane associated large macromolecular/amyloid-like structures to sediment along with crude membranes (Liu *et al*, 2017). Ubiquitylated MLKL emerged and accumulated within the whole cell lysate and the crude membrane fraction from 90 minutes post-stimulus onwards, coinciding precisely with the appearance of phosphorylated MLKL and onset of cell death (**Fig. 3A & Supp. Fig. 1A**). On the other hand, the predominant species of MLKL in the cytosol was non-phosphorylated and non-ubiquitylated (**Fig. 3A)**. These data suggest that MLKL ubiquitylation occurs in the crude membrane fraction but not in the cytosol.

**Figure 3.**
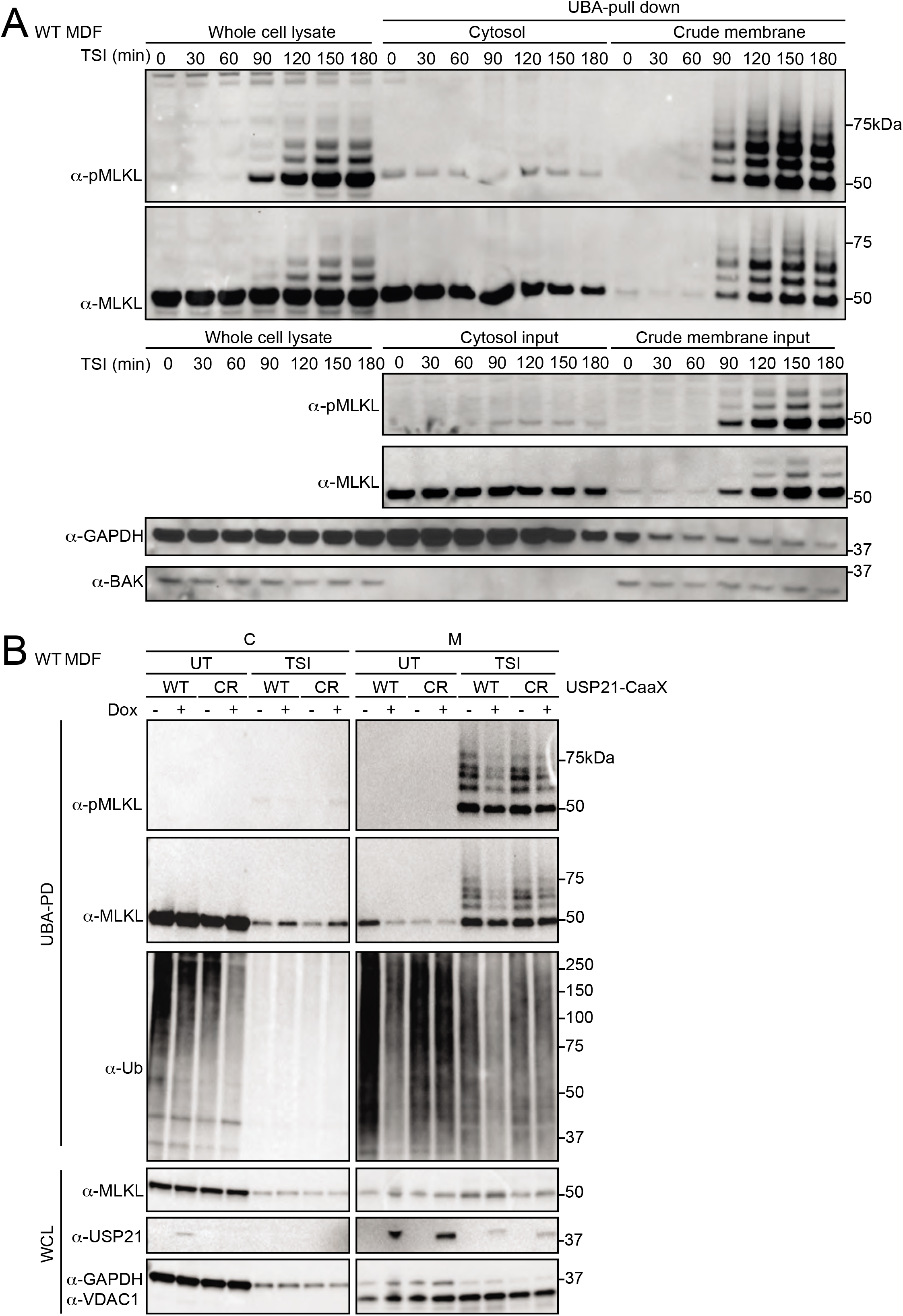
MLKL ubiquitylation accumulates in the crude membrane fraction and can be digested by USP21 located on biological membranes. A WT MDFs were treated with TSI for indicated time. Cells were fractionated into cytosol and crude membrane parts, followed by UBA-pull down. All fractions were analysed by Western blot and probed with antibodies as indicated. Representative of three independent experiments. B WT USP21-CaaX and USP21^C221R^-CaaX (CR) were inducibly expressed in WT MDFs by doxycycline for 5 hrs. Ubiquitylated proteins were enriched followed by TSI stimulation and cellular fractionation. All fractions were analysed by Western blot and probed with antibodies as indicated. Representative of three independent experiments.

To explore further where ubiquitylated MLKL is located we took advantage of the fact that ubiquitylated MLKL was digested by USP21 (**Fig. 2B**). We therefore fused a CaaX motif tag (C=cysteine, a=aliphatic amino acid, X=terminal residue) to the C-terminus of the catalytic domain USP21 to localise it to the plasma membrane (Hancock *et al*, 1991). We chose ‘CVLQ’ to mimic the CaaX motif found at the C-terminal end of another DUB, USP32 (Sapmaz *et al*, 2019), and also generated a catalytically-dead USP21 mutant version (USP21^C221R^) as a control, also fused with CaaX motif. This mutant, like the previously published mutant USP21^C221A^, lacks the ability to remove ubiquitin (Morrow *et al*, 2018; Ye *et al.,* 2011; Ye *et al*, 2012). Using these stably and inducibly expressed constructs we evaluated TSI-mediated MLKL ubiquitylation with or without USP21-CaaX function. Taking into account the amount of MLKL in the crude membrane fraction it was clear that while it did not completely denude MLKL of ubiquitin, the WT USP21-CaaX fusion did substantially reduce the amount of ubiquitylated MLKL, as well as generally affecting the levels of ubiquitylated proteins (**Fig. 3B**). Taken together these data suggest that ubiquitylated MLKL is localised to the plasma membrane.

### Necroptosis induced MLKL ubiquitylation is driven by its oligomerization

In order to kill, MLKL oligomerizes, translocates to, and permeabilises membranes. To dissect further the drivers of MLKL ubiquitylation, we generated a series of MLKL mutants that were defective in one or more of these essential steps. Previously, we found that alanine replacement of the surface exposed residues R105 and D106 in the four-helix bundle (4HB) domain of mouse MLKL prevents it from oligomerising, translocating and inducing cell death (Hildebrand *et al.,* 2014; Tanzer *et al.,* 2016). In the context of full length MLKL, we found that these mutations prevent formation of a high molecular weight complex (complex II) in the ‘crude membrane’ fraction following necroptotic stimulation by TSQ (**Fig. 4A**). Although R105A/D106A MLKL was phosphorylated following this necroptotic stimulus, it did not exhibit the distinct TSI-induced ubiquitylation observed for WT MLKL (**Fig. 4B**). This supports the idea that MLKL only undergoes ubiquitylation post activation by RIPK3 and suggests that MLKL does not undergo ubiquitylation without oligomerization.

**Figure 4.**
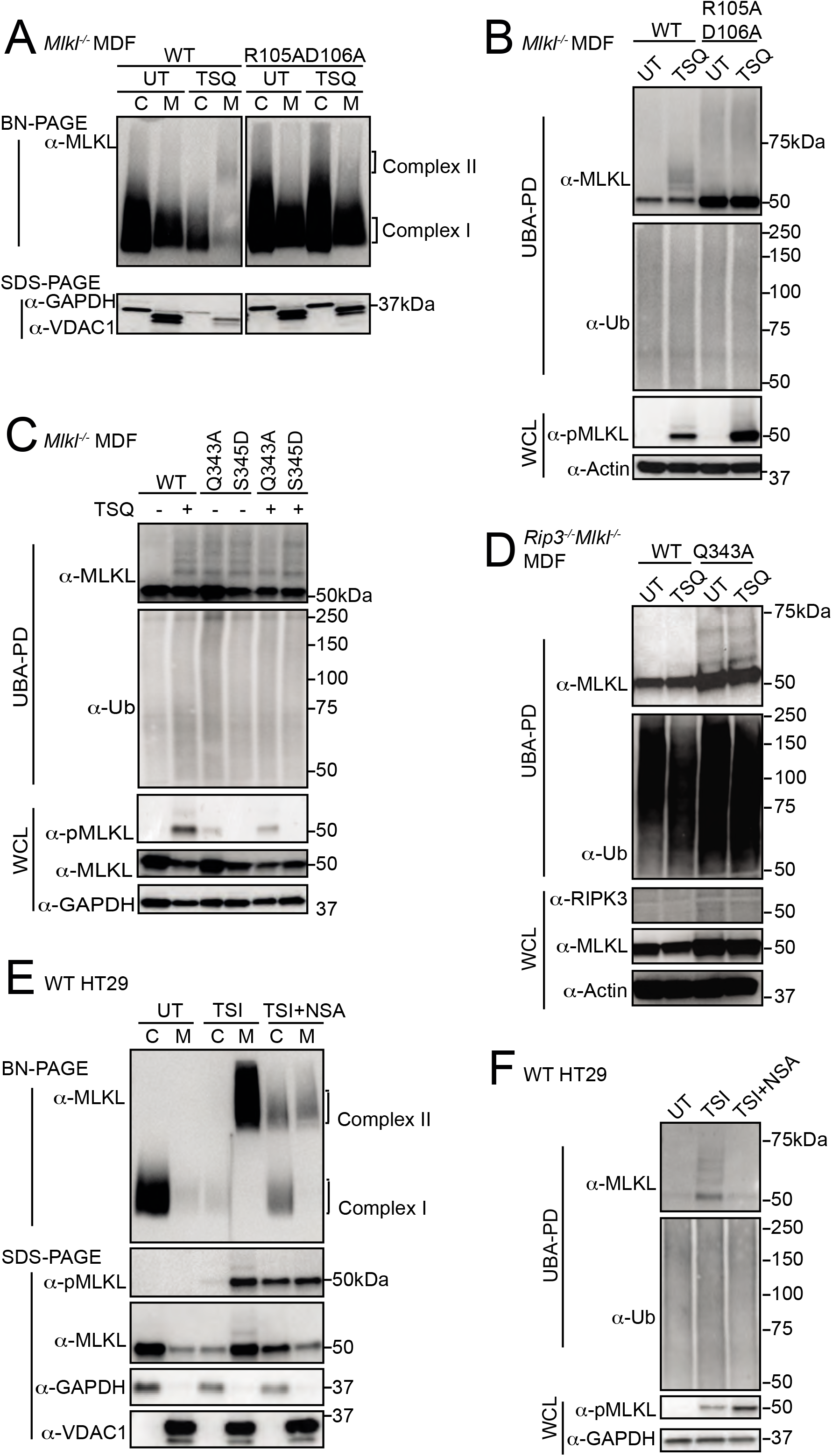
MLKL oligomerization drives its necroptosis specific ubiquitylation. A WT and R105AD106A mutant MLKL were inducibly expressed in *M1k1^-/-^* MDFs by doxycycline, at the same time cells were untreated (UT) or treated with for 6 hrs. Cells were fractionated into cytosol (C) and crude membrane (M). Fractions were analysed by BN-or SDS-PAGE, Western blot and probed with the indicated antibodies. Representative of three independent experiments. B Cell lysates from (A) were subjected to UBA-pull down and analysed as described above. C WT, Q343A and S345D mutant MLKL were inducibly expressed in *M1k1^-/-^* MDFs by doxycycline, at the same time cells were treated ± TSQ for 16 hrs, followed by UBA-pulldown. Representative of three independent experiments. D WT and Q343A mutant MLKL were inducibly expressed in *Ripk3^-/-^ M1k1^-/-^* MDFs by doxycycline, at the same time cells were treated ± TSQ for 16 hrs, followed by UBA-pulldown. Representative of three independent experiments. E HT29 cells were stimulated with TSI, ± NSA (500 nM), or left untreated (UT) for 16 hrs, followed by cellular fractionation. Fractions were analysed by BN-or SDS-PAGE, Western blot and probed with the indicated antibodies. Representative of three independent experiments. F Cell lysates from (E) were subjected to UBA-pulldown and analysed as described above.

The MLKL mutants, Q343A (which perturbs a hydrogen bond with K219 in the ATP binding motif VTIK) and S345D (phospho-mimetic), are auto-activated forms of murine MLKL (Murphy *et al.,* 2013). We induced expression of Q343A or S345D mutant MLKL in *M1k1^-/-^* and *Ripk3^-/-^ M1k1^-/-^* MDFs for 16 hrs where, as predicted, they killed cells independently of an extrinsic necroptotic stimulus (**Supp. Fig. 4B**). As expected, induction of necroptosis in *Mlkl* cells reconstituted with wild type MLKL resulted in MLKL ubiquitylation (**Fig. 4C**). However, activated MLKL mutants expressed in the same *M1k1^-/-^* MDFs were ubiquitylated independently of a necroptotic stimulus and to an extent comparable to wildtype MLKL induced to undergo necroptosis (**Fig. 4C**). Furthermore, Q343A MLKL became ubiquitylated in *Ripk3^-/-^ M1k1_-/-_* MDFs and therefore independently of any upstream activation (**Fig. 4D**). We conclude that while MLKL ubiquitylation correlates with its oligomerization and phosphorylation, direct RIPK3 association is not required for MLKL ubiquitylation in murine cells.

Necrosulfonamide (NSA) is an inhibitor of human MLKL by forming a covalent bound with Cys86 that only exists in human MLKL but not mouse MLKL (Liu *et al.,* 2017). Consistent with a recent report (Murai *et al*, 2018; Samson *et al*, 2020), we found that while it did not inhibit the phosphorylation of MLKL or the formation of high molecular weight MLKL-containing complex, large proportions of this MLKL species remained in the cytosolic (0.025% digitonin soluble) fraction (C) of cells (**Fig. 4E**). Consistent with the reduction in MLKL oligomer in the crude membrane (0.025% digitonin insoluble) fraction (M), NSA also reduced MLKL ubiquitylation (**Fig. 4F**).

### Ubiquitylated MLKL undergoes proteasome and lysosome dependent turnover

Unlike the R105A/D106A MLKL mutant, E109A/E110A MLKL mutant and N-terminal FLAG-tagged MLKL (N-FLAG MLKL) are able to form higher order oligomers in crude membrane fractions following necroptotic stimulation, yet are nevertheless unable to induce necroptosis (Hildebrand *et al.,* 2014; Tanzer *et al.,* 2016) (**Fig. 5, B, Supp. Fig. 5A, B**). These mutants therefore allowed us to examine whether ubiquitylation was a consequence of necroptosis. Upon induction of necroptosis, E109A/E110A MLKL was more heavily phosphorylated and ubiquitylated than wildtype MLKL (**Fig. 5C**). Similarly, N-FLAG MLKL was also phosphorylated and ubiquitylated within 2 hrs of TSI treatment, although these modified MLKL species diminished over time (**Fig. 5D**).

**Figure 5.**
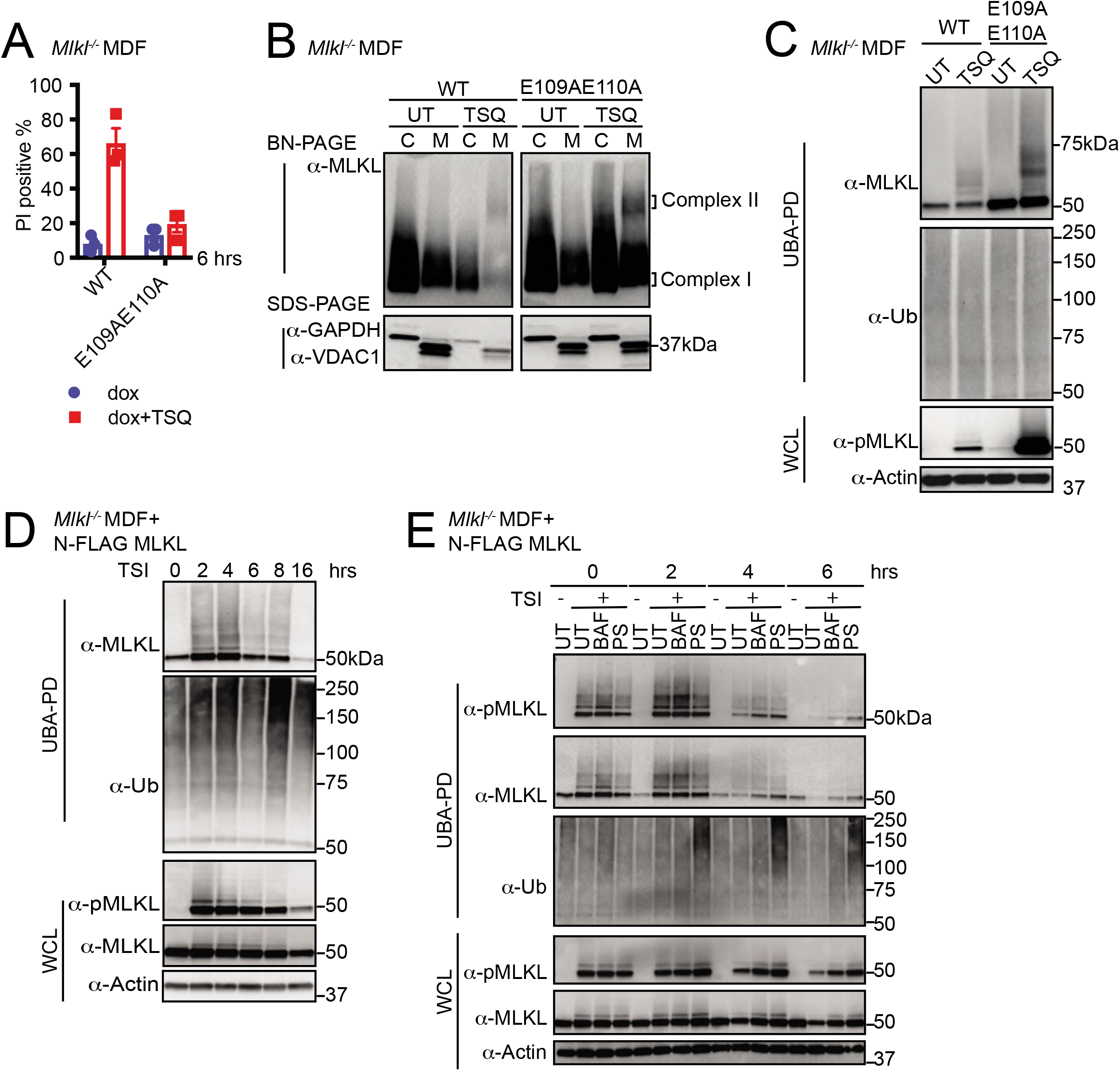
MLKL-ubiquitylation does not lead to cell death, but correlates with the turnover of activated MLKL. A WT and E109AE110A mutant MLKL were inducibly expressed in *M1k1^-/-^* MDFs by doxycycline, cells were untreated (UT) or treated with TSQ for 6 hrs (please note that this same control was used in Figure 4A). Cell death was measured by PI staining based on flow cytometry. Data are plotted as mean ± SEM of three independent experiments. B Cellular fractions from (A) were analysed by Western blot from BN-PAGE or SDS-PAGE using antibodies as indicated. Representative of three independent experiments. C Cell lysates from (A) were subjected to UBA-pull down and analysed as described above. D N-FLAG MLKL were inducibly expressed in *M1k1^-/-^* MDFs by doxycycline overnight, and cells were treated with TSI for indicated time, followed by UBA-pull down. Representative of three independent experiments. E N-FLAG MLKL were inducibly expressed in *M1k1^-/-^* MDFs by doxycycline for 16 hrs. Cells were stimulated ± TSI for 3 hrs after withdrawal of doxycycline. Then TSI medium was removed and replaced to medium containing inhibitors Bafilomycin A1 (BAF), PS341 (PS) or left untreated (UT). IDN-6556 was added to all conditions to block apoptosis. Cells were collected 0, 2, 4, 6 hrs after medium replacement, followed by UBA-pull down. Representative of three independent experiments.

The cellular turnover of active forms of MLKL has been observed previously by our group and by others (Gong *et al*, 2017; Hildebrand *et al.,* 2020; Yoon *et al*, 2017; Zargarian *et al*, 2017). Therefore, we examined ubiquitylation of N-FLAG MLKL in MDFs over a time course following a TSI pulse. N-FLAG MLKL is phosphorylated, oligomerizes and accumulates in the 0.025% insoluble cell fraction without killing *M1k1^-/-^* MDFs following necroptotic stimulation. This later feature facilitates the study of these ubiquitylated MLKL species and their cellular turnover *in situ* using proteasome and lysosome inhibitors as previously described for unmodified MLKL (Hildebrand et al, 2020). We found that after 4 hrs TSI stimulation, levels of ubiquitylated and phosphorylated N-FLAG MLKL dramatically decreased. This decrease can be delayed by either the lysosome inhibitor bafilomycin or the proteasome inhibitor PS341 (**Fig. 5E**). This suggests that MLKL ubiquitylation correlates with its proteasome and lysosome dependent turnover.

### MLKL ubiquitylation antagonises necroptosis

We sought to identify the precise amino acids on MLKL that were modified by ubiquitin. We enriched for activated and ubiquitylated N-FLAG MLKL from the ‘crude membrane’ fraction of MDFs by performing a FLAG tag affinity purification. Eluted fractions from FLAG affinity beads were analysed by mass spectrometry, which identified Gly-Gly conjugates on lysine residues K9, K51, K69 and K77 (**Supp. Fig. 6A**). All four are located in the 4HB domain of MLKL (**Supp. Fig. 6B**) which suggests a role for ubiquitylation in regulating MLKL-mediated cell death. We did not identify any additional signs of ubiquitin modification on the brace or pseudokinase region of MLKL using this experimental system.

We mutated the four lysines to arginine and inducibly expressed the “4KR” mutant in *M1k1^-/-^* MDFs to explore their role. Surprisingly, the 4KR MLKL was phosphorylated and ubiquitylated in a similar manner to wild type MLKL following TSI treatment (**Fig. 6B**). There was a reduction in the ability of this mutant to kill cells in response to TSI treatment (**Supp. Fig. 6C**), but this is seemingly unrelated to its ubiquitylation and more likely due to modification of conserved residues in the cytotoxic domain of MLKL. These results highlight the difficulties in mutational analyses when studying ubiquitylation and rather than mutate any of the other 38 lysines in MLKL we tried a different approach.

**Figure 6.**
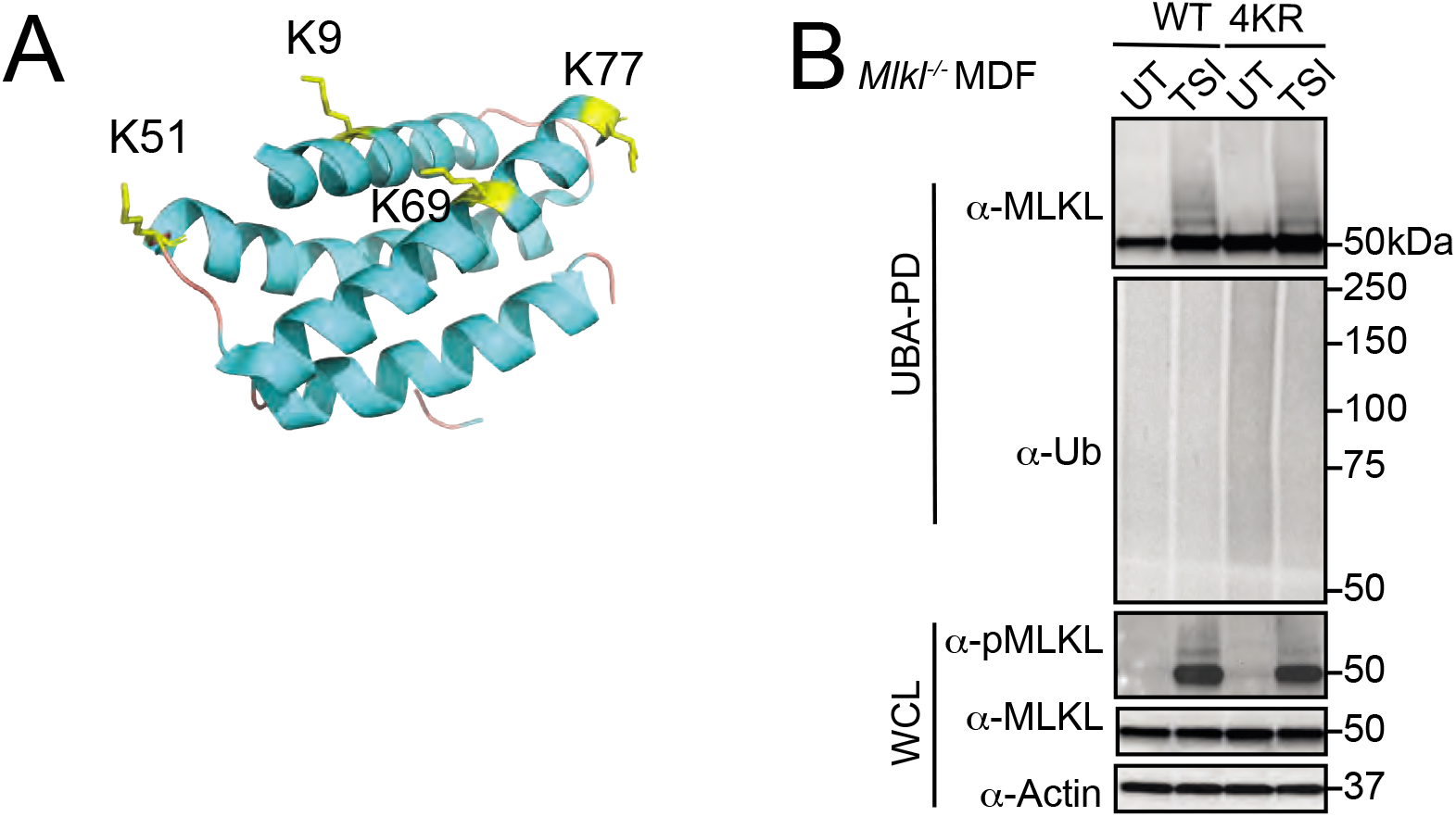
Simultaneous arginine replacement of 4 ubiquitylation sites on the mMLKL 4HB domain does not prevent necroptosis-induced ubiquitylation. A Cartoon of the N-terminal region (residues 1-180) of mouse MLKL (PDB accession 4BTF;(Murphy *et al.,* 2013)) showing the four lysine residues identified from MS analysis as yellow sticks. B WT and 4KR mutant MLKL were inducibly expressed in *M1k1^-/-^* MDFs by doxycycline for 6 hrs and cells were untreated (UT) or treated with TSI, followed by UBA-pull down. Representative of three independent experiments.

We had observed that the deubiquitylating enzyme USP21 removed all ubiquitin from MLKL *in vitro* (**Fig. 2B**). We therefore hypothesised that fusing the catalytic domain of USP21 to MLKL would remove TSI induced ubiquitylation from MLKL. We fused it to the C-terminal end of MLKL, because, as with the FLAG tag, N-terminal fusions affect the ability of MLKL to kill (**Supp. Fig. 5A**). We fused the catalytic domain of USP21 to the C-terminus of both human and mouse MLKL (**Fig. 7A**) and expressed the fusion proteins in *MLKL^-/-^* HT29 and *M1k1^-/-^* MDF cells respectively, to determine, firstly, whether it still retained cytotoxic activity and, secondly, whether it would be resistant to necroptosis induced ubiquitylation. To specifically control for the loss of ubiquitylation, we also generated MLKL fused to a catalytically-dead USP21 mutant (USP21^C221R^; **Fig. 7A**), as mentioned in **Fig. 3B**. Both the MLKL-USP21 and MLKL-USP21^C221R^ fusion proteins were exogenously expressed and the cells were stimulated with TSI for varying amounts of time. Ubiquitylated proteins were enriched *via* UBA-pulldown. Like murine MLKL^WT^, the catalytically inactive control, MLKL-USP21^C221R^, when expressed in *M1k1^-/-^* MDFs, showed high MW laddering indicative of ubiquitylation and the ubiquitylation was enhanced following TSI stimulation (**Fig. 7B**).

**Figure 7.**
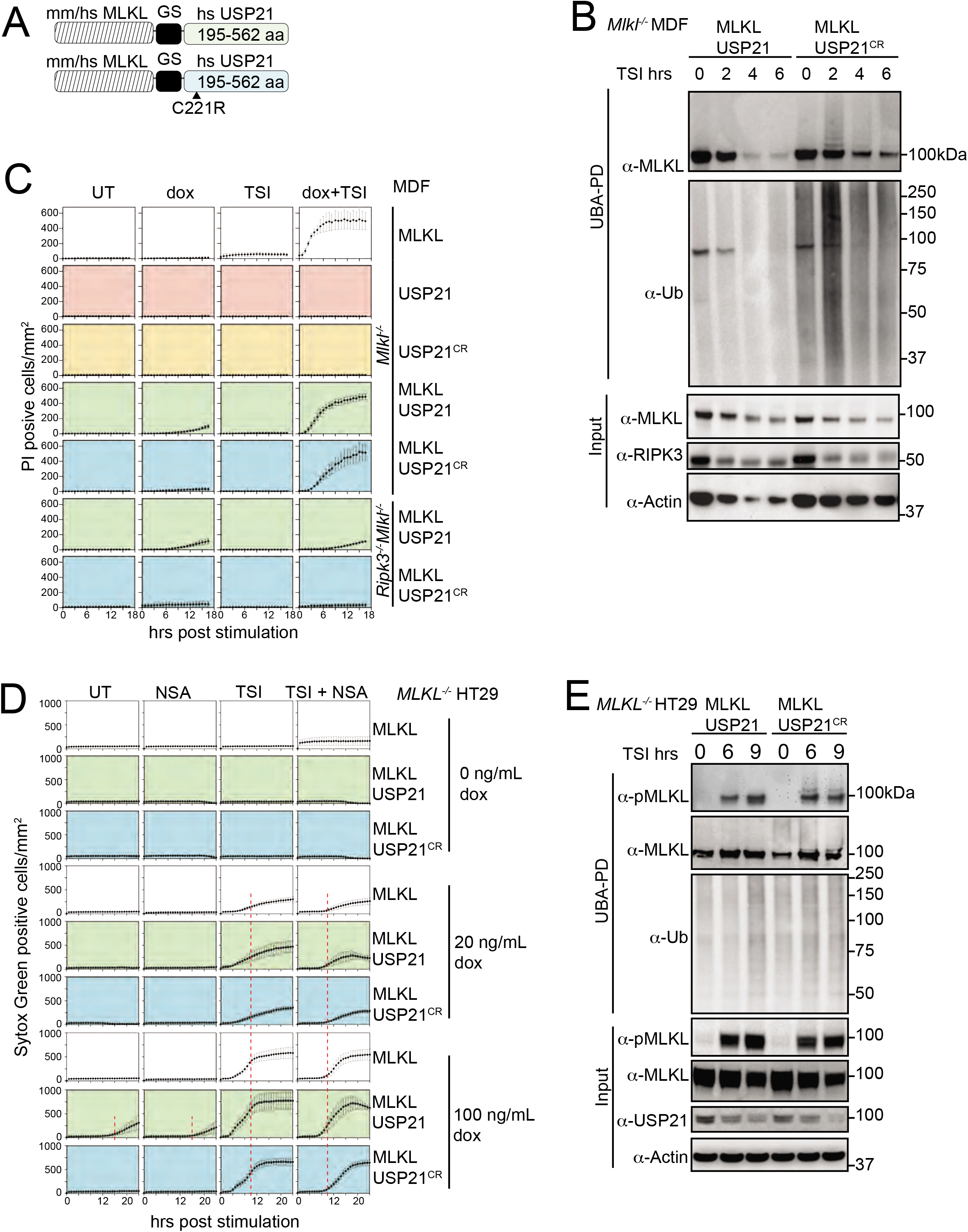
MLKL ubiquitylation antagonises necroptosis. A Schematic of the MLKL-USP21 and USP21 catalytic dead mutant proteins. B Mouse MLKL-USP21 and MLKL-USP21^C221R^ were inducibly expressed in *M1k1^-^* MDFs by doxycycline (10 ng/mL) for 6 hrs with addition of a necroptotic stimulus (TSI) for indicated time, followed by UBA-pulldown. Representative of three independent experiments. C *M1k1^-/-^* MDFs and *Ripk3^-/-^ M1k1^-/-^* MDFs stably transfected with constructs encoding MLKL, USP21, USP21^C221R^, MLKL-USP21 and MLKL-USP21^C221R^ were treated with doxycycline, TSI or in combination. Propidium iodide positive cells were quantified in real time by IncuCyte live cell imaging. Representative of three independent experiments. D *MLKL^-/-^* HT29 cells stably transfected with constructs encoding human MLKL-USP21 and MLKL-USP21^C221R^, were treated with doxycycline, NSA (1 μM), TSI or combinations thereof (added simultaneously). Sytox Green positive cells were quantified in real time by live cell imaging. Representative of 6 independent experiments. (A red dashed line is shown to highlight the delay in death kinetics upon treatment with NSA). E Human MLKL-USP21 and MLKL-USP21^C221R^ were inducibly expressed in *MLKL^-/-^* HT29 cells by doxycycline (10 ng/mL) for 16 hrs and then cells were treated with TSI for the indicated time course, followed by UBA-pulldown. Representative of three independent experiments.

These high MW MLKL-USP21^C221R^ species were reduced and collapsed into non-ubiquitylated form by digestion of recombinant USP21 **(Supp. Fig 7C)**. In contrast, the ubiquitin laddering was not evident for the MLKL-USP21^WT^ fusion (**Fig. 7B**). MLKL-USP21 and MLKL-USP21^C221R^ re-constituted sensitivity to TSI stimulation in *M1k1^-/-^* MDFs with only modestly delayed kinetics when compared to wildtype MLKL (**Fig. 7C**). However, unlike wildtype MLKL or MLKL-USP21^C221R^, MLKL-USP21 could induce cell death independent of TSI stimulation (**Fig. 7C**), even when expressed in *Ripk3^-/-^ M1k1^-/-^* MDFs (**Fig. 7C**), without being phosphorylated (see Input fractions in **Supp. Fig. 7C**).

We made similar observations for human MLKL-USP21 fusions expressed in *MLKL^-/-^* HT29 cells. Human MLKL-USP21^C221R^, but not MLKL-USP21 fusions, became ubiquitylated following TSI stimulation (**Fig. 7E**). Furthermore, like human MLKL, human MLKL-USP21 and MLKL-USP21^C221R^ fusions were able to reconstitute the capacity of *MLKL^-/-^* HT29 cells to undergo TSI induced necroptosis when their expression was induced by doxycycline (**Fig. 7D)**, while USP21 controls did not (**Supp. Fig. 7B)**. For the first 10 hrs of expression and activation, NSA blocked TSI-induced cell death for all MLKL species, which was then overcome by overexpression. MLKL-USP21 also exhibited TSI-independent cell death, occurring from 16 hrs, which can also be delayed by NSA. (Fig. 7D). While phosphorylated MLKL was not detected by that time (**Fig. 7E**). These data indicate that ubiquitylation of MLKL is not required for necroptotic cell death, but rather acts as an important kinetic regulator of the necroptosis pathway following oligomerization and membrane association.

## Discussion

MLKL undergoes ubiquitylation during necroptosis. In this study, we identified distinctive ubiquitylation of both mouse and human MLKL that could be induced by a range of necroptotic stimuli. This signature ubiquitin ladder was essentially resistant to cleavage by a host of DUBs. Only USP21, the DUB that can remove all ubiquitin modifications, including mono-ubiquitylation, was able to deubiquitylate MLKL. Therefore, we propose that MLKL is mono-ubiquitylated at multiple sites. Mono-ubiquitylation is typically associated with endosome-lysosome trafficking (Haglund *et al*, 2003; Mosesson & Yarden, 2006) and is less likely to play a scaffolding role or result in proteasomal degradation (Oh *et al*, 2018; Wilkinson *et al*, 1995).

Full activation of MLKL’s killing activity is a multi-step and protracted process, the choreography of which has not yet been fully deduced. Nevertheless there is a broad consensus, based on analysis of MLKL mutants and a range of different techniques, that phosphorylation by RIPK3 leads to MLKL oligomerization, translocation to biological membranes and subsequent membrane permeabilization (Petrie *et al*, 2020; Petrie *et al.,* 2018; Samson *et al.,* 2020; Tanzer *et al.,* 2016). Recent single-cell imaging approaches examining human MLKL show that phosphorylated MLKL clusters in cytoplasmic vesicles together with RIPK1. These vesicles are actively transported to the plasma membrane, where they begin to coalesce in hotspots, some hours prior to membrane lysis (Samson *et al.,* 2020). Consistent with data showing that NSA does not prevent MLKL phosphorylation or initial oligomerization, but does block higher order oligomerization (Liu *et al.,* 2017)), NSA appears to prevent MLKL clustering (Samson *et al.,* 2020). To define where and when ubiquitylation of MLKL occurred we fractionated cells into cytosolic and crude membrane fractions, and only detected ubiquitylated MLKL in the 0.025% digitonin insoluble (crude membrane) fraction. We also examined a number of different MLKL mutants that we had previously characterised for their ability to form oligomers, translocate to the crude membrane fraction and promote necroptosis. The MLKL mutants, Q343A and S345D, which are able to form 0.025% digitonin insoluble high molecular weight oligomers and induce cell death without necroptotic stimulation (Murphy *et al.,* 2013; Tanzer *et al*, 2015), were ubiquitylated independently of RIPK3 or a necroptotic TSQ stimulus. On the other hand, the R105A/D106A mutant that is unable to form high molecular weight oligomers on BN-PAGE did not become ubiquitylated following a necroptotic stimulus. This implies that MLKL ubiquitylation occurs after MLKL oligomerization and does not specifically require RIPK3 or upstream signalling.

MLKL has been variously reported to traffic to the nucleus (Weber *et al*, 2018; Yoon *et al*, 2016), lysosomes (Fan *et al*, 2019; Wang *et al*, 2014; Yoon *et al.,* 2017), mitochondria (Wang *et al*, 2012) and the plasma membrane (the most likely location) (Cai *et al.,* 2014; Chen *et al.,* 2014; Hildebrand *et al.,* 2014; Samson *et al.,* 2020) following necroptotic activation. Crude membrane fractionation enriches membrane and insoluble protein aggregates, therefore in an attempt to determine more precisely where MLKL ubiquitylation occurred, we generated a plasma membrane-targeted, CaaX-fused USP21 (Wright & Philips, 2006). Because ubiquitin modification plays a role in the initial, plasma membrane located, TNF/TNFR1 induced necroptotic signalling, we anticipated that expressing USP21-CaaX might affect induction of necroptosis. However necroptosis was induced in cells expressing a non-targeted USP21 or a catalytically inactive USP21^C221R^-CaaX to equal levels as in USP21-CaaX cells, despite the fact that USP21-CaaX was clearly active and reduced total ubiquitin levels. Consistent with MLKL being targeted to the plasma membrane, MLKL was less ubiquitylated in USP21-CaaX cells than in cells expressing USP21^C221R^-CaaX following a necroptotic stimulus. MLKL ubiquitylation was, however, not completely prevented by USP21-CaaX expression leaving open several interpretations of this data viz: not all MLKL is ubiquitylated at the plasma membrane; or USP21-CaaX was unable to access all MLKL in the membrane; or MLKL is ubiquitylated prior to reaching the plasma membrane and can only be deubiquitylated once it arrives there. Therefore while we cannot exclude that ubiquitylated MLKL sediments as part of large amyloid-like polymers or large cytoskeletal structures in the crude membrane fraction without being directly associated with a biological membrane (Liu *et al.,* 2017), our data suggest that at least some ubiquitylated MLKL is accessible to a plasma membrane localised DUB.

As previously observed (Murai *et al.,* 2018), we found that in the presence of NSA, the high molecular weight human MLKL complex that is normally found exclusively in the crude membrane fraction was now also observed in the cytosolic fraction. Combined with other observations (Samson *et al.,* 2020), this indicates that NSA interferes with MLKL membrane association. Since NSA treatment also inhibited MLKL-ubiquitylation, this suggests that membrane association is required for MLKL ubiquitylation, and that the relevant E3 ligase is also membrane associated.

Neither N-terminally FLAG tagged MLKL nor the E109A/E110A mutant MLKL are capable of killing cells, although both are phosphorylated and form 0.025% digitonin-insoluble high molecular weight oligomers in a similar manner to wild type MLKL following induction of necroptosis. Since both are also ubiquitylated we can conclude that ubiquitylation is not sufficient to cause necroptosis. To more precisely define the role of MLKL ubiquitylation during necroptosis we used mass spectrometry to identify ubiquitylated lysines in mouse MLKL. We found four such lysines in the 4HB domain, but none in the pseudokinase domain. Only one of these four, K77, is conserved in human MLKL, however this is not particularly surprising because homology of charged residues in MLKL is not well conserved between mouse and human (**Supp. Fig. 6B**).

Mutation of these lysines did not prevent either MLKL induced killing or MLKL ubiquitylation. The persistence of ubiquitin modification is a relatively common scenario when trying to define the role of ubiquitylation by mutating lysine. This is because E3 ligase mediated ubiquitylation is rarely tightly restricted to a motif and can therefore be promiscuous in target modification (Petroski & Deshaies, 2003; Wu *et al*, 2003). This means that if a favoured lysine in the target is mutated, the E3 ligase may nevertheless ubiquitylate another lysine. We therefore tried a more innovative approach. Since we had shown that the DUB USP21 was able to remove necroptosis induced ubiquitylation of MLKL *in vitro,* we generated an MLKL C-terminal USP21 fusion that we predicted would be constitutively deubiquitylated. A similar approach fusing the K63 specific DUB AMSH to EGFR has been used to investigate EGFR degradation (Huang *et al*, 2013), however it was not clear whether a DUB that removes all ubiquitin would be as well tolerated by cells. Stable inducible expression of USP21 alone was not toxic to cells over 24 hours and fusion of the wildtype or catalytically-dead USP21 did not markedly affect MLKL’s cytotoxic activity following a TSI death stimulus. Furthermore, catalytically-active USP21 fused to MLKL prevented TSI induced ubiquitylation of MLKL. Interestingly, in both human and mouse cells, loss of MLKL ubiquitylation also allowed MLKL to kill cells without a necroptotic stimulus, albeit less potently when compared with such a stimulus. This further supports the idea that ubiquitylation of MLKL is an important ‘insurance policy’ against low level activation of MLKL by other cellular kinases or low level spontaneous transition to the active conformation. Furthermore our results indicate that the approach of directly fusing a DUB, even a pan DUB like USP21, to a protein of interest may be a widely applicable technique to evaluate the role of ubiquitin modification in other systems.

Other mechanisms have been proposed for MLKL turnover post-activation. The ESCRT-III machinery was proposed to mediate plasma membrane shedding alongside active MLKL during necroptosis (Gong *et al.,* 2017), and cells were reported to release active MLKL containing vesicles via endosomal trafficking (Fan *et al.,* 2019; Yoon *et al.,* 2017; Zargarian *et al.,* 2017). It has been proposed that this turnover helps set a threshold of activated MLKL required for necroptosis and that it delays the onset of cell death to allow production of essential cytokines and Damage-associated molecular patterns (DAMPs) and an inflammatory response (Vandenabeele *et al*, 2017). Another possibility is that the prolonged multi-step path from MLKL phosphorylation to membrane permeabilization and cell death allows multiple points of regulation that ensure that a cell only commits to necroptosis under precisely defined conditions (Dovey *et al*, 2018; Hildebrand *et al.,* 2020; Jacobsen *et al*, 2016; Samson *et al.,* 2020). For this to work, the cell must have mechanisms to deactivate already activated MLKL, and the ubiquitylation of MLKL may serve as one such mechanism. Our results, showing that MLKL-USP21, but not a wild type MLKL expressed to similar levels, kills cells in the absence of a necroptotic stimulus suggest that MLKL may be continually translocating to membranes and turned over by MLKL-ubiquitylation. Like the rapid degradation of inflammatory cytokine mRNAs (Lacey *et al*, 2015; Menon & Gaestel, 2018) although energetically costly, this might allow a more rapid response to a pathogen than an on/off mechanism focused solely on initiation and may also provide another mechanism to detect and act on attempts by pathogens to interfere with this anti-pathogen response.

## Materials and Methods

### Compounds and cytokines

Human TNF-Fc made in-house, Smac-mimetics Compound A (Tetralogic), IDN-6556 (Tetralogic), Q-VD-OPh (R & D Systems), Lipopolysaccharide (LPS, Sigma), Poly I:C (Sigma), Fas Ligand (a gift from Lorraine O’Reilly, WEHI), doxycycline (Sigma), Necrostatin-1 (Nec-1, Sigma), GSK872 (a gift from Anaxis Pty, Ltd.), coumermycin (Sigma), Necrosulfonamide (NSA, Merck Millipore), Propidium iodide (Sigma), Sytox Green (Thermo Fisher), puromycin (Thermo Fisher), N-ethylmaleimide (NEM, Sigma), deubiquitylase (DUBs) made in house (Hospenthal *et al.,* 2015), complete protease inhibitor cocktail (Roche), Bafilomycin A1 (BAF, Enzo), PS341 (Sigma).

### Antibodies

Mouse phospho-MLKL (Ser345) rabbit monoclonal ERP9515(2) (Abcam)

Mouse phospho-MLKL (Ser345) rabbit monoclonal (D6E3G) (Cell Signalling Technology) MLKL rat monoclonal 3H1 (available from Millipore MABC604; generated in-house) ß-actin mouse monoclonal AC-15 (Sigma A-1978)

BAK NT rabbit polyclonal #06-536 (EMD Millipore)

GAPDH rabbit monoclonal 14C10 (Cell Signalling Technology)

Pan-ubiquitin mouse monoclonal #3939 (Cell Signalling Technology) Mouse/human RIPK1 mouse monoclonal #610458 (BD transduction Laboratories)

Mouse RIPK3 rabbit polyclonal PSC-2283-c100 (Axxora (Pro Sci))

Human RIPK3 rat monoclonal 1H2 (generated in-house)(Petrie *et al*, 2019b) Human phospho-MLKL (Ser358) rabbit monoclonal ab 187091 (Abcam) VDAC1 rabbit polyclonal AB10527 (Millipore)

Human Usp21 rabbit polyclonal 17856-I-AP (Proteintech) HRP-conjugated secondary antibodies

### Cell lines immortalization and transfection

Cells were cultured in DMEM+8% FCS at 37°C. HT29 cells were a kind gift from Mark Hampton. MDFs were isolated from tails of mice bearing different genotype, and immortalized by SV40 large T antigen via lentivirus transduction. MLKL mutant constructs were generated as described previously ((Hildebrand *et al.,* 2014; Moujalled *et al.,* 2014; Murphy *et al.,* 2013)). All oligonucleotides for PCR and mutagenesis were synthesized by IDT. Constructs encoding USP21 catalytic domain with GS linker were made by GenScript (Nanjing, CN). Genes encoding WT mouse MLKL and PCR-derived mutants were cloned into pF TRE3G PGK puro, a puromycin-selectable, doxycycline-inducible vector as previously described (and kindly supplied by Toru Okamoto) (Hildebrand *et al.,* 2014; Moujalled *et al.,* 2014; Murphy *et al.,* 2013; Tanzer *et al.,* 2016). Lentiviruses were generated in HEK293T cells before infection of target cells. Puromycin (5 μg/mL) was added for selection and maintenance of lines stably transduced with lentivirus.

For BMDMs, femora and tibiae were collected from WT and *Tnf^-/-^* mice at 6 weeks old, cut and flushed by PBS. Bone marrow were incubated in Petri dishes with DMEM supplemented with 10% FCS and 20% conditioned L929 medium for 7 days. Cells were fed with fresh medium after 3 days of plating.

### Concentrations of stimuli and inhibitors

TNF (100 ng/mL), Smac-mimetic Compound A (500 nM), IDN-6556 (5 μM), Q-VD-OPh (5 μM), doxycycline (20 ng/mL), Necrostatin-1 (50 μM), GSK872 (5 mM), LPS (50 ng/mL), Fas ligand (6.4 ng/mL), Poly I:C (1 mg/mL), coumermycin (700 nM), Bafilomycin (5 μM), PS341 (50 nM), MG132 (200nM), Chloroquine (50 μM), NH4Cl (2 mM), Ca-074 Me (20 μM), TLCK (100 μM), AEBSF (100 μM) and concentrations otherwise indicated.

### Cell death measured by flow cytometry

Cells were plated at 50,000/well in 24-well plates before treatment, collected by trypsin and spun down, resuspended and stained with PI (Propidium Iodide, 1 μg/mL) in PBS buffer, and quantified by Flow cytometry.

### Cell death measured by live cell imaging

MDFs were plates at 10,000/well or 15,000/well (96-well plates) and HT29s were plated at 40,000/well (48-well plates) or 15,000/well (96-well plates) and allowed to settle for 4 hrs (MDFs) and 24 hrs (HT29s) respectively. Cells were stimulated as indicated in media containing Propidium Iodide (PI, 200 ng/mL) or Sytox Green (500nM, ThermoFisher Scientific) and imaged at 45 min or 1 hr intervals using an IncuCyte S3 Live cell imager. Numbers of PI or Sytox positive cells were quantified and plotted using IncuCyte software.

### Cellular fractionation and Blue Native-PAGE

Cells were collected by scraping, spun down and washed in pre-chilled PBS. MELB buffer (20 mM 4-(2-Hydroxyethyl) piperazine-1-ethanesulfonic acid, (HEPES) pH=7.5, 100 mM sucrose, 2.5 mM MgCl_2_, 100 mM KCl) containing 0.025% digitonin was used to permeabilise cells to extract the cytosol fraction. Non-soluble part was further solubilized by 1% digitonin buffer. (Liu *et al.,* 2018; Schagger & von Jagow, 1991) Both fractions were separated by Bis-Tris Native PAGE, and transferred to PVDF membrane. After transfer the membrane was then destained (50% methanol and 25% acetic acid) and denatured (in 6M guanidine HCl, 5 mM β-ME) to maximise epitope exposure.

### UBA-pull down assay and UbiCRest

Recombinant GST-UBA fusion protein (Hjerpe *et al*, 2009) purified from *E. coli* in house was prebound to glutathione sepharose beads (10 μL/condition). Cells were lysed in DISC buffer (30 mM Tris-HCl, pH 7.5, 150 mM NaCl, 10% glycerol) containing 1% Triton X-100 with 10 mM NEM and protease inhibitor. Cleared lysate was then coupled with beads overnight. SDS-PAGE sample loading buffer was used to elute the UBA-pull down fraction. For UbiCrest, washed beads were then incubated with DUBs at previously established concentrations (Hospenthal *et al.,* 2015) and incubated at 37°C for the indicated times. Digestion product was eluted in SDS sample buffer for Western blot analysis. Beads elution fractions were generated by removing the digestion product from the beads and following washing.

### FLAG-tagged protein immunoaffinity precipitation and Mass Spectrometry sample preparation

N-FLAG WT mouse MLKL was inducibly expressed in *M1k1^-/-^* MDFs by doxycycline overnight and stimulated with TSI for 3 hrs. Cells were collected by scraping and permeabilised by 0.025% digitonin in MELB buffer together with NEM (10 mM) and protease inhibitors. The crude membrane fraction was pelleted by centrifugation and dissolved in 1% SDS in DISC buffer. Cleared lysate of crude membrane was then applied to M2-FLAG beads for affinity precipitation. FLAG tagged protein on beads was eluted by heating at 56°C in 0.5% SDS solution. Eluted material was subjected to tryptic digestion using the FASP method (Wisniewski *et al*, 2009). Peptides were lyophilised using CentriVap (Labconco) prior to reconstituting in 80 μl 0.1% FA/2% acetonitrile (ACN). Peptide mixtures (1 μl) were analysed by nanoflow reversed-phase liquid chromatography tandem mass spectrometry (LC-MS/MS) on an M-Class HPLC (Waters) coupled to a Q-Exactive Orbitrap mass spectrometer (Thermo Fisher). Peptide mixtures were loaded in buffer A (0.1% formic acid, 2% acetonitrile, Milli-Q water), and separated by reverse-phase chromatography using C18 fused silica column (packed emitter, I.D. 75 μm, O.D. 360 μm x 25 cm length, IonOpticks, Australia) using flow rates and data-dependent methods as previously described (Delconte *et al*, 2016; Kedzierski *et al*, 2017). Raw files consisting of high-resolution MS/MS spectra were processed with MaxQuant (version 1.5.8.3) for feature detection and protein identification using the Andromeda search engine (Cox *et al*, 2011). Extracted peak lists were searched against the UniProtKB/Swiss-Prot *Mus musculus* database (October 2016) and a separate reverse decoy database to empirically assess the false discovery rate (FDR) using strict trypsin specificity allowing up to 2 missed cleavages. The minimum required peptide length was set to 7 amino acids. The mass tolerance for precursor ions and fragment ions were 20 ppm and 0.5 Da, respectively. The search included variable modifications of oxidation (methionine), amino-terminal acetylation, carbamidomethyl (cysteine), GlyGly or ubiquitylation (lysine), phosphorylation (serine, threonine or tyrosine) and N-ethylmaleimide (cysteine).

## Data availability

The raw mass spectrometric data and the MaxQuant result files have been deposited to the ProteomeXchange Consortium via the PRIDE (Perez-Riverol *et al*, 2019) partner repository with the dataset identifier: PXD015537.

**Username:** **Password:** f2WTYnC4

## Acknowledgements

We would like to thank Jiami Han, Yueyuan Li and the WEHI mouse facility for technical assistance. This work was funded by NHMRC grants 1025594 (JS), 1046984 (JS) and 1105023 (JS and JMH) and fellowships 1172929 (JMM), 1058190 (JS), 1107149 (JS) and 110574 (JS) and was made possible through Victorian State Government Operational Infrastructure Support and Australian Government NHMRC IRIISS (9000587).

## Author Contribution

ZL, LD and KSA designed and performed experiments, and analysed data. ZL, JMH and JS analysed the data and wrote the manuscript. DK conceived the DUB fusion experiment and provided reagents. All authors read and commented on the manuscript.

## Conflict of interest statement

SNY, AB, UN, CF, SEG, JMM, JMH and JS contribute to, or have contributed to, a project with Anaxis Pty Ltd to develop necroptosis inhibitors.

## Abbreviations

T: TNF
S: Smac-mimetics Compound A
I: IDN-6556
Q: Q-VD-OPh
WCL: whole cell lysate
UBA-PD: pull down fractions
C: cytosolic fraction
M: crude membrane fraction
WT: wildtype
CR: C221R
PI: Propidium Iodide
Nec-1: Necrostatin-1
Ub: ubiquitin and others as indicated elsewhere
TS, SI: TSI
TSQ: are used in combination, as apoptotic or necroptotic stimuli

---

**Supplementary Figure 1.**
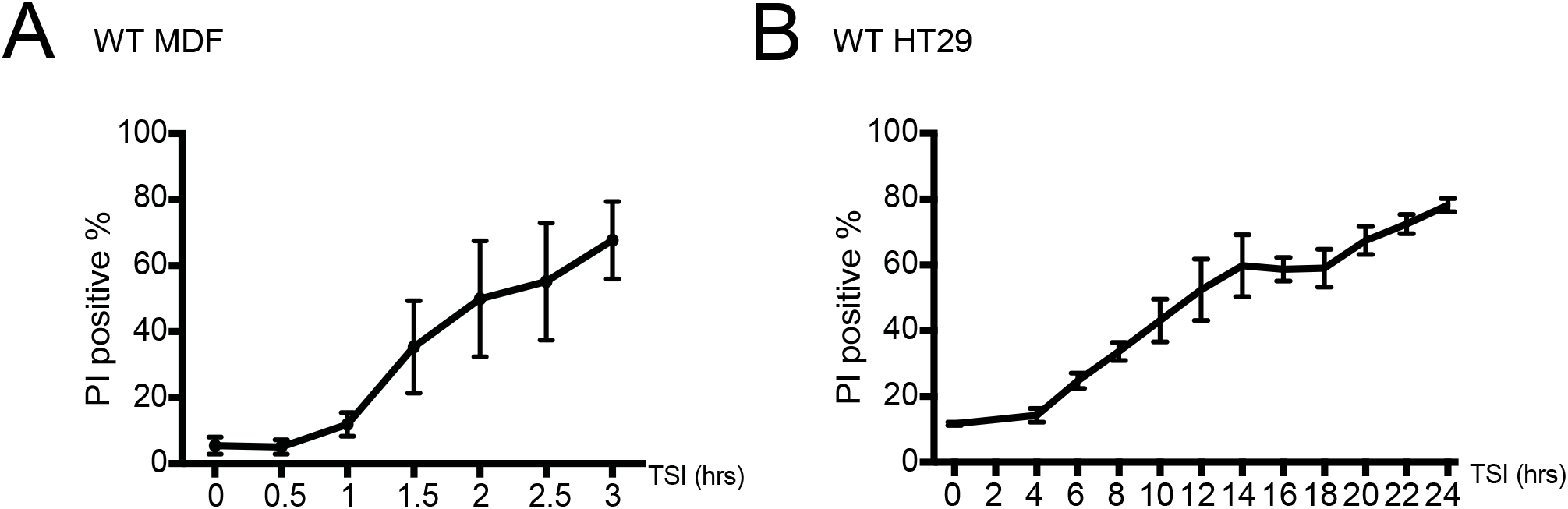
Cell death time course of MDFs and HT29 cells following necroptotic stimulation. MDFs (A) and HT29 (B) cells were treated with TSI. Cell death was measured by PI staining and flow cytometry. Data are plotted as mean ± SEM of three independent experiments.

**Supplementary Figure 3.**
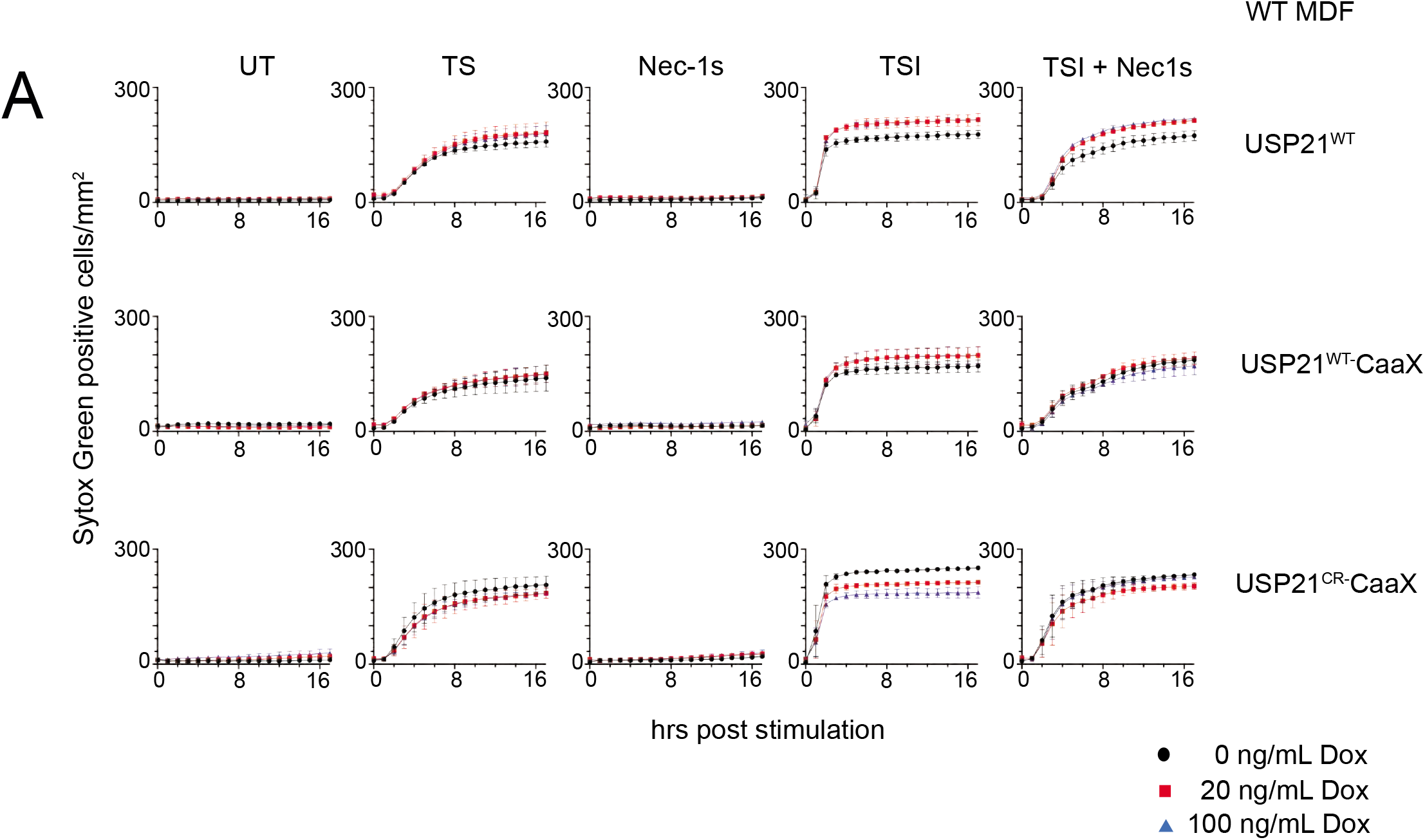
USP21-CaaX expression does not alter the kinetics of TNF induced apoptosis or necroptosis in MDFs. A WT USP21, USP21-CaaX and USP21^C221R^-CaaX were inducibly expressed in WT MDFs by doxycycline. TS and Nec-1 were used to control for apoptotic signalling. Sytox Green positive cells were quantified in real time by live cell imaging. Representative of 2 independent experiments.

**Supplementary Figure 4.**
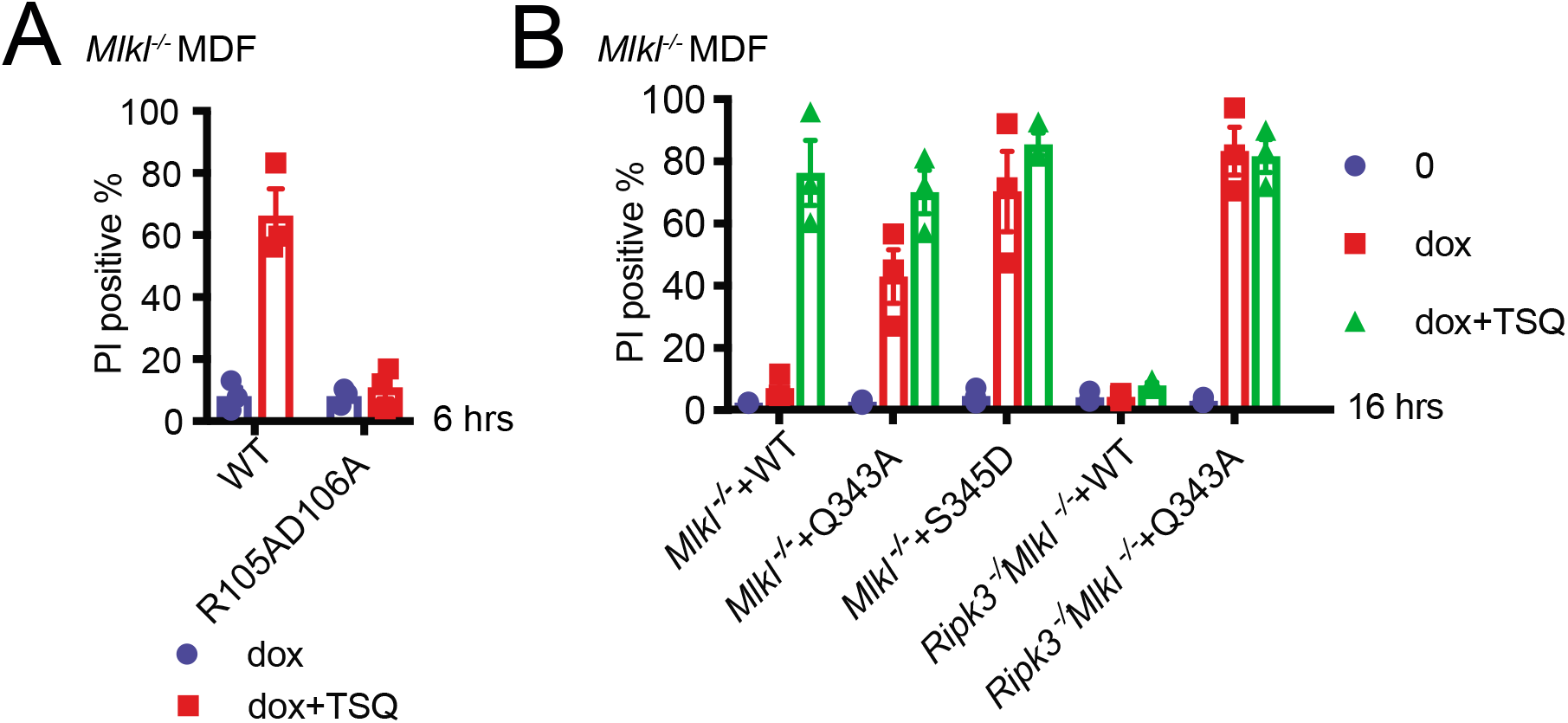
MLKL oligomerization drives its necroptosis specific ubiquitylation. A Cell death of samples from **Fig. 4A** was measured by PI staining based on flow cytometry. Data are plotted as mean ± SEM of three independent experiments. B Cell death of samples from **Fig. 4C, D** were analysed as in (A).

**Supplementary Figure 5.**
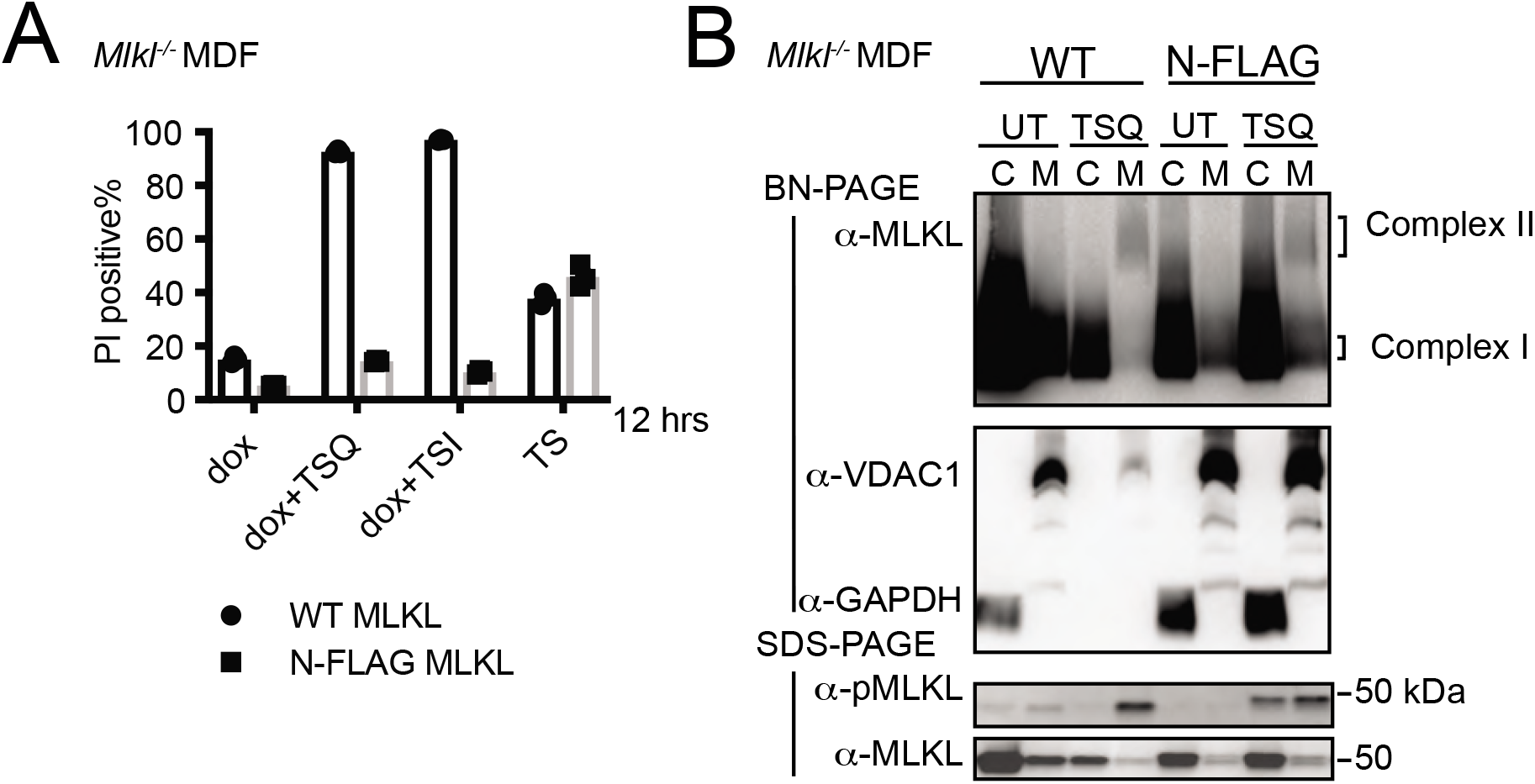
N-FLAG MLKL behaves like WT MLKL but does not induce cell death following necroptotic stimulation. A WT MLKL and N-FLAG MLKL were inducibly expressed in *M1k1^-/-^* MDFs by doxycycline for 12 hrs and cells were treated with the TSI or TSQ necroptotic stimuli. TS was included to control for responding to apoptosis signalling. Cell death was measured by PI staining based on flow cytometry. Data are plotted as mean ± SEM of three independent experiments. B WT MLKL and N-FLAG MLKL were inducibly expressed in *M1k1^-/-^* MDFs by doxycycline for 6 hrs, cells were untreated (UT) or treated with TSQ. Cellular fractions were analysed by Western blot from BN-PAGE or SDS-PAGE using antibodies as indicated. Representative of three independent experiments.

**Supplementary Figure 6.**
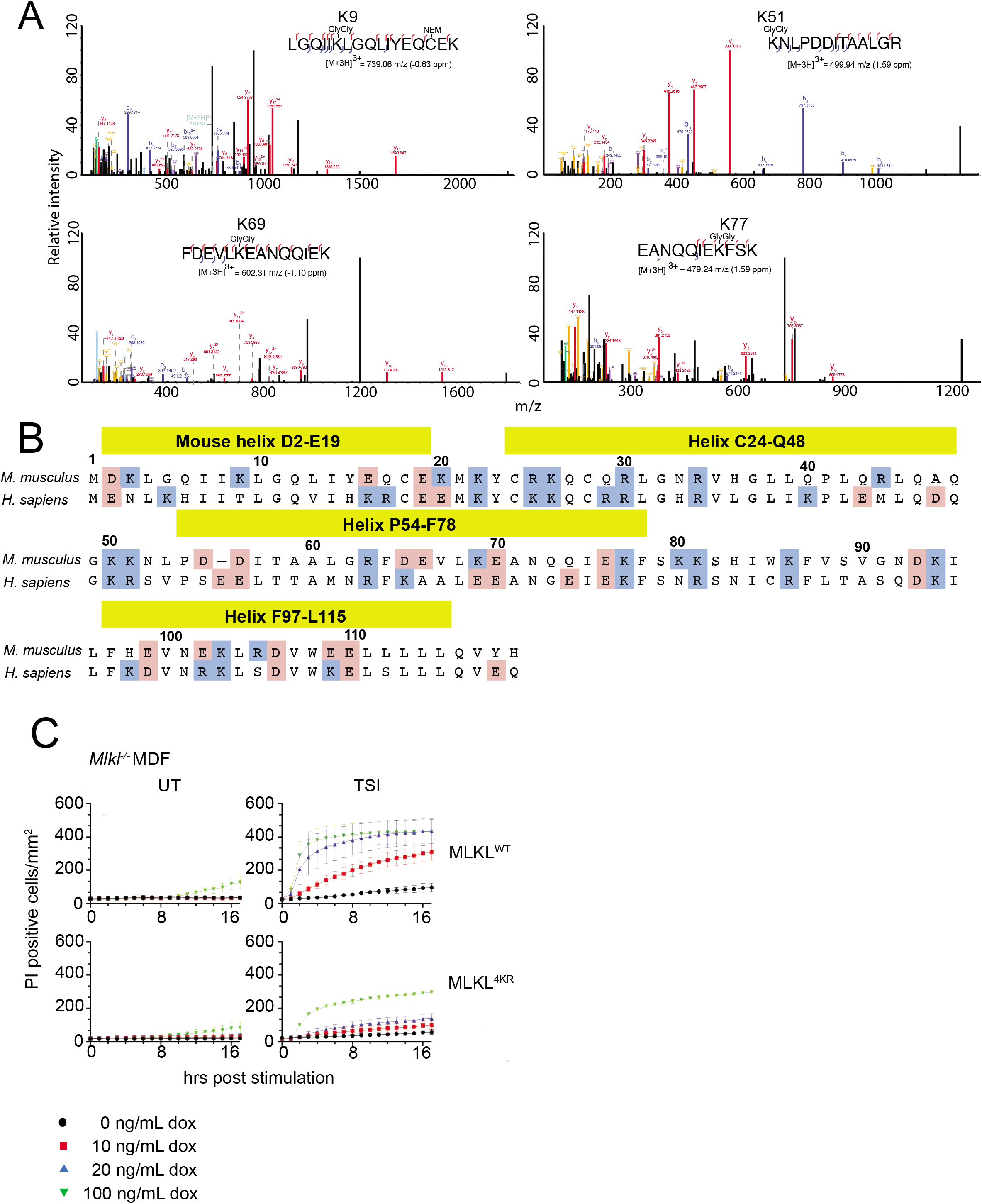
Simultaneous arginine replacement of 4 ubiquitylation sites on the mouse MLKL 4HB domain does not prevent necroptosis-induced ubiquitylation. A MS spectra were manually validated to confirm the identification of four Gly-Gly sites on activated MLKL. B Alignment of mouse and human MLKL N-terminal domain. Positively charged residues are labelled in blue and negatively charged residues are labelled in pink. C WT and 4KR mutant MLKL were inducibly expressed in *M1k1^-/-^* MDFs by doxycycline and cells were treated ±TSI (added simultaneously) for 4 hrs. Sytox Green positive cells were quantified in real time by IncuCyte S3 live cell imaging. Representative of 3 independent experiments.

**Supplementary Figure 7.**
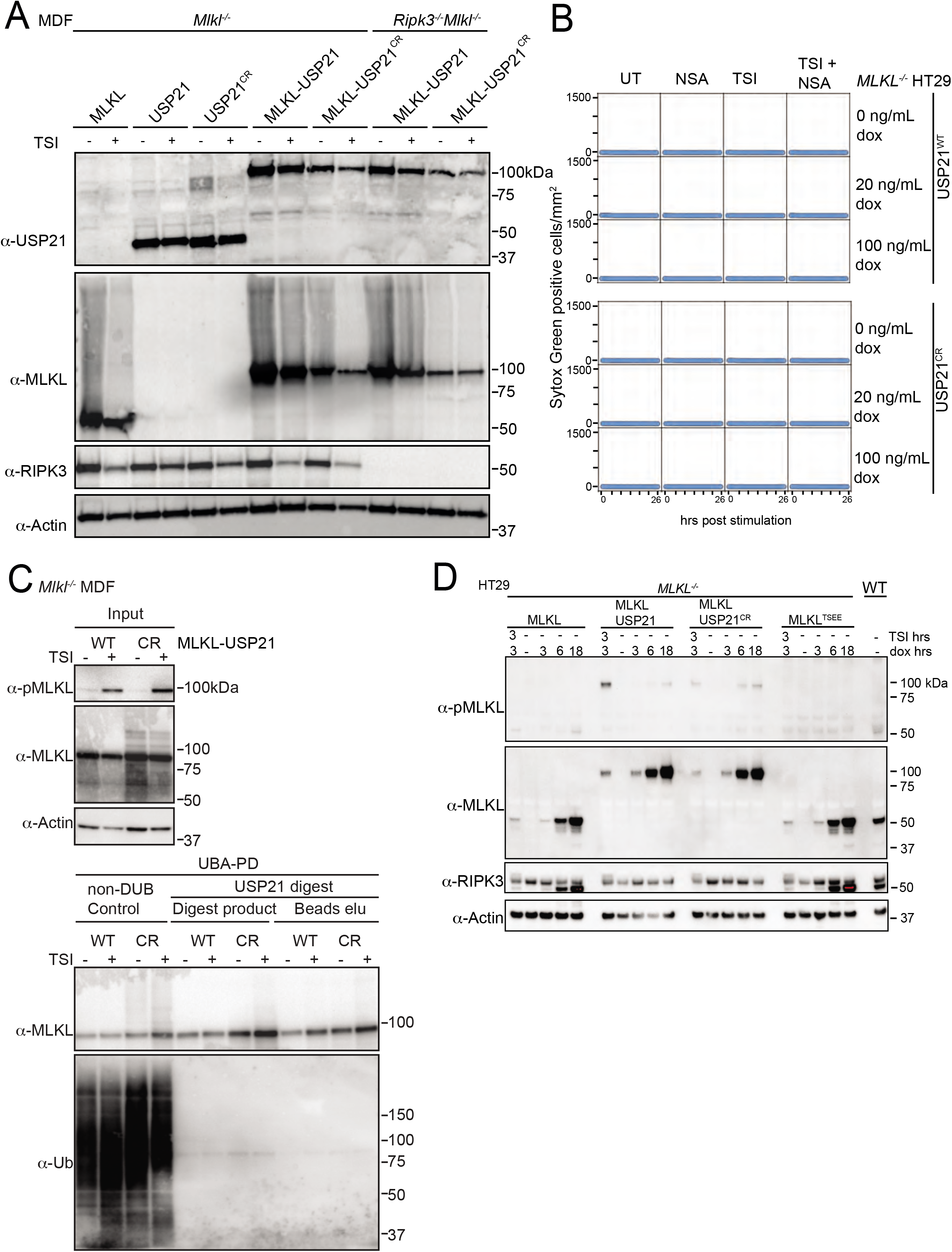
MLKL ubiquitylation antagonises necroptosis. **A** Mouse MLKL, human USP21 and mouse MLKL-human USP21 fusions were inducibly expressed in *Ripk3^-/-^ M1k1^-/-^* MDFs following doxycycline addition (20 ng/mL) for 6 hours ± TSI. Representative of 2 independent experiments. **B** *MLKL^-/-^* HT29 cells stably transfected with doxycycline inducible constructs encoding human USP21 and human USP21^C221R^ were treated with doxycycline, NSA (1 μM), TSI or combinations thereof (added simultaneously). Sytox Green positive cells were quantified in real time by live cell imaging. Representative of 2 independent experiments. **C** Mouse MLKL-USP21 and MLKL-USP21^C221R^ were inducibly expressed in *M1k1^-/-^* MDFs by doxycycline (10 ng/mL) for 8 hrs with addition of a necroptotic stimulus (TSI) for 3hrs, followed by UBA-pulldown and USP21 digestion. Antibody (D6E3G, Cell Signaling Technology) was used here to detect MLKL phosphorylation. Representative of 2 independent experiments. **D** *MLKL^-/-^* HT29 cells were stably transfected with indicated doxycycline inducible *MLKL* alleles (here phosphor-mimic human MLKL mutant T357E/S358E indicated as MLKL^TSEE^) and treated with doxycycline (100 ng/mL) ± TSI (added simultaneously). Representative of 3 independent experiments.

